# Quantifying the linkages between California sea lion (*Zalophus californianus*) strandings and particulate domoic acid concentrations at piers across Southern California

**DOI:** 10.1101/2023.08.15.549094

**Authors:** Jayme Smith, Jacob A. Cram, Malena Berndt, Vanessa Hoard, Dana Shultz, Alissa C. Deming

## Abstract

Domoic acid producing blooms of the diatom genus *Pseudo-nitzschia* are pervasive in coastal environments globally. Domoic acid, a neurotoxin, accumulates via trophic transfer into marine food webs and are often associated with mass marine mammal mortality and stranding events. In Southern California, California sea lions (*Zalophus californiaus*) are an indicator species for food web impacts of domoic acid because they are abundant secondary consumers, sensitive to domoic acid intoxication, and are actively monitored by stranding networks. However, domoic acid exposure may occur a distance from where a sea lion ultimately strands. This spatiotemporal variation complicates coupling domoic acid observations in water to strandings. Therefore, we sought to quantify whether monitoring data from four pier sites across the region, covering nearly 700 km of coastline from 2015-2019, could be used to predict adult and subadult sea lion strandings along the 68 km Orange County coastline surveyed by the Pacific Marine Mammal Center. We found that increased sea lion strandings were often observed just prior to an increase in particulate domoic acid at the piers, confirming that clusters of subadult and adult sea lion strandings with clinical signs of domoic acid intoxication a serve as indicators of bloom events. In addition, domoic acid concentrations at Stearns Wharf, nearly 200 km from stranding locations, best predicted increased total sea lion strandings, and strandings of sea lions with domoic acid intoxication symptoms. Particulate domoic acid concentrations greater than 0.05 μg/L at Stearns Wharf led to a detectable increase in stranding probability in Orange County, and concentrations over 0.25 μg/L resulted in a nearly 1.6-fold increase in stranding probabilities for a given week.

## Introduction

Harmful algal blooms (HABs) are a common water quality issue in many coastal and inland waterbodies globally and cause widespread impacts on human and animal health, recreation, fisheries and aquaculture (Broadwater et al., 2018; Moore et al., 2020; Wells et al., 2020; Anderson et al., 2021). One of the most pervasive HAB forming organisms, particularly within the Southern California, is the diatom genus *Pseudo-nitzschia,* which contains over a dozen species capable of producing the water-soluble neurotoxin domoic acid (Bates et al., 2018). Domoic acid accumulates via trophic transfer into pelagic and benthic food webs, resulting in significant risks to both human and ecosystem health (Bates et al., 1989; Bejarano et al., 2008b; Lefebvre et al., 2017). Domoic acid exposure causes sickness and mass mortality events in a variety of marine animals including sea lions, sea otters, whales and seabirds (Lefebvre et al., 2002a; Kvitek et al., 2008; Torres De La Riva et al., 2009; Gibble et al., 2021; Moriarty et al., 2021).

The first event in which domoic acid intoxication of marine animals was conclusively linked to a bloom of *Pseudo-nitzschia* occurred in 1998 in Monterey Bay, California. During this event, over 70 California sea lions (*Zalophus californiaus*) were treated for domoic acid intoxication, 48 of which died (Scholin et al., 2000). Domoic acid was detected not only in the blood and urine of the affected sea lions but also was detected in sardines, anchovies, and phytoplankton in the area (Scholin et al., 2000). Since this event, domoic acid related strandings of marine mammals have continued to be observed for more than two decades along the California coastline, impacting hundreds of marine animals (Gulland et al., 2002; Greig et al., 2005; Bejarano et al., 2008a, 2008b; Goldstein et al., 2008).

Marine mammals are good sentinels of coastal ecosystem health due to their high trophic status, distinctive life histories, and long term residence in coastal environments, which are traits that make them responsive to ecosystem changes (Rice and Rochet, 2005; Bossart, 2011). Within Southern California, California sea lions are an important indicator species due to their permanent residence in the region, abundance, and their upper trophic level status (Melin et al., 2012). California sea lions are also commonly rescued and rehabilitated by the California Marine Mammal Stranding Network (Greig et al., 2005). California sea lions strand for a variety of reasons including malnutrition, infectious diseases, human interactions (ie: gunshot wounds, fishing line entanglement), natural and anthropogenic changes in the environment (storms, climatic cycles, climate change), and exposure to algal toxins such as domoic acid (Greig et al., 2005; Simeone et al., 2015; Gulland et al., 2022).

Domoic acid is a water-soluble chemical analog of L-glutamate and is an agonist of glutamate receptors. It causes massive neuronal activation, leading to seizures, neuronal injury, and neuronal death over time (Silvagni et al., 2005). Domoic acid intoxication is most often diagnosed by clinical signs combined with a known bloom occurring in the region of the strandings. Diagnostic testing for domoic acid in feces, urine, stomach contents, as well as other fluids and tissues are available, but not always pursued due to the pathopneumonic signs associated with acute domoic acid intoxication. California sea lions can have different clinical presentations to domoic acid exposure based on whether the animal has had an acute intoxication or has developed chronic symptoms in response to multiple exposures over time, although the progression from acute to chronic is not well understood. California sea lions exposed acutely to domoic acid typically display a common suite of neurological symptoms including seizures, postictal behavior, head weaving, ataxia, lethargy, and abnormal scratching (Scholin et al., 2000; Gulland et al., 2002). Other symptoms include abortion, premature parturition of pups and death (Brodie et al., 2006). Acute cases are reported to strand in spatiotemporal clusters (Bejarano et al., 2008a). Domoic acid epileptic disease, also referred to as chronic epileptic syndrome in sea lions, is thought to be caused by one or more sublethal exposure events resulting in permanent brain damage (Ramsdell and Gulland, 2014). Sea lions with this syndrome show behavioral changes, intermittent seizures (suffering from seizures two weeks apart or after two weeks in rehabilitation) associated with hippocampal atrophy (Goldstein et al., 2008). These animals are often asymptomatic between seizures and strand individually, unlike acute domoic acid stranding events that present in stranding clusters with dozens of sea lions display DA intoxication clinical signs within a small geographic region. Adult females are most frequently associated with domoic acid related strandings in California, although all sexes and age classes of sea lions are susceptible to intoxication (Bejarano et al., 2008a).

The relationship between California sea lion stranding events and the presence of domoic acid in the environment is complex for multiple reasons. One major factor is the temporal and spatial variability of domoic acid in the environment, particularly during bloom events. The production of domoic acid by *Pseudo-nitzschia* is dynamic and varies based on environmental conditions, which can also influence bloom formation and duration (Trainer et al., 2020). Blooms are also heterogeneous spatially. The level of toxin present in *Pseudo-nitzschia* cells can vary across both horizontal and vertical gradients along the coast (Seegers et al., 2015; Bowers et al., 2018). The amount of toxin that is produced during a bloom event is also dependent on the *Pseudo-nitzschia* species present, since not all species of *Pseudo-nitzschia* produce toxins, and the toxin production rates of toxigenic species can vary considerably across species and strains (Trainer et al., 2012; Bowers et al., 2018). Another complicating factor is that domoic acid can enter the food web via multiple mechanisms. Domoic acid can enter both pelagic (Lefebvre et al., 1999, 2002b) and benthic food webs (Vigilant and Silver, 2007; Smith et al., 2021), and marine mammal exposure is then mediated by the consumption of contaminated prey items that are typically primary or secondary consumers. Recent work has also shown that sea lion prey organisms can differentially accumulate domoic acid (Bernstein et al., 2021), further complicating the linkages between the presence of domoic acid in water samples and impacts on upper trophic levels.

California sea lions are a useful indicator species for identifying pelagic food web impacts of domoic acid producing blooms in the region. Previous studies along the California coastline have highlighted correlations between the presence of either domoic acid or *Pseudo-nitzschia* cells in the environment and sea lion stranding events. Significant, but weak, correlations between a monthly time series of sea lion strandings of all age classes and abundances of two different toxigenic *Pseudo-nitzschia* species and particulate domoic acid (pDA) concentrations were observed between 1998-2007 in Monterey Bay, California (Bargu et al., 2010). In Southern California, a significant, multi-species marine mammal stranding event was caused by a large domoic acid producing bloom in 2002. Comparisons between stranding reports and *Pseudo-nitzschia* relative abundances, a semiquantitative measure of *Pseudo-nitzschia* abundance in relation to other phytoplankton, for the year of 2002 indicated significant correlations between strandings of multiple marine mammal species and increased relative abundances of *Pseudo-nitzschia* both immediately and with lags up to six weeks following peaks in *Pseudo-nitzschia* relative abundances (Torres De La Riva et al., 2009).

Here, we aim to identify a quantitative link between observations of domoic acid in water samples and marine mammal strandings to build towards the ability to predict negative ecosystem impacts from bloom events. This study considered particulate domoic acid data collected routinely from 2015-2019 at four pier stations across the region and California sea lions stranding data from the same period collected along a 68 km stretch of the Southern California coastline by the Pacific Marine Mammal Center (PMMC). The goals of this study were to: 1) model the relationship between pier based domoic acid data and sea lion strandings 2) identify which pier(s) were most strongly linked with stranding events, and 3) to identify thresholds of domoic acid concentrations that are associated with an elevated risk of stranding events.

## Materials and Methods

### Pier Based, Offshore, and Subsurface Water Sampling

Ambient harmful algal bloom monitoring for domoic acid concentrations and other related parameters has been conducted on a weekly basis since 2008 within Southern California as a part of the California Harmful Algal Bloom Monitoring and Alert Program (HABMAP; Kudela et al., 2015). Particulate domoic acid (pDA) concentrations from January 2015 to December 2019 were monitored at four pier locations within the Southern California (**Figure 1**): Scripps Pier in San Diego (n = 260), Newport Beach Pier on the San Pedro Shelf (n = 256), Santa Monica Pier in the Santa Monica Bay (n = 258), and Stearns Wharf along the Santa Barbara Channel (n = 258). The collection and analysis of pDA samples are described in Seubert et al. (2013). Briefly, surface water samples were collected weekly, typically on Mondays, at each HABMAP pier location. Subsamples were collected in duplicate for pDA analysis via gentle vacuum filtration of 188 - 200 mL of sample water onto glass fiber filters. Filters were stored at -20 °C in the dark until analyzed. Filters were extracted in 3 mL of 10% methanol, sonicated for 15 seconds and centrifuged for 15 minutes at 4,000 rpm. The supernatant was analyzed via Mercury Science, Inc., Domoic Acid Enzyme-Linked ImmunoSorbant Assay (ELISA: Mercury Science, Durham, NC) according to the methods described in Litaker et al. (2008). The detection limit for pDA is 0.02 μg/L.

**Figure 1.**
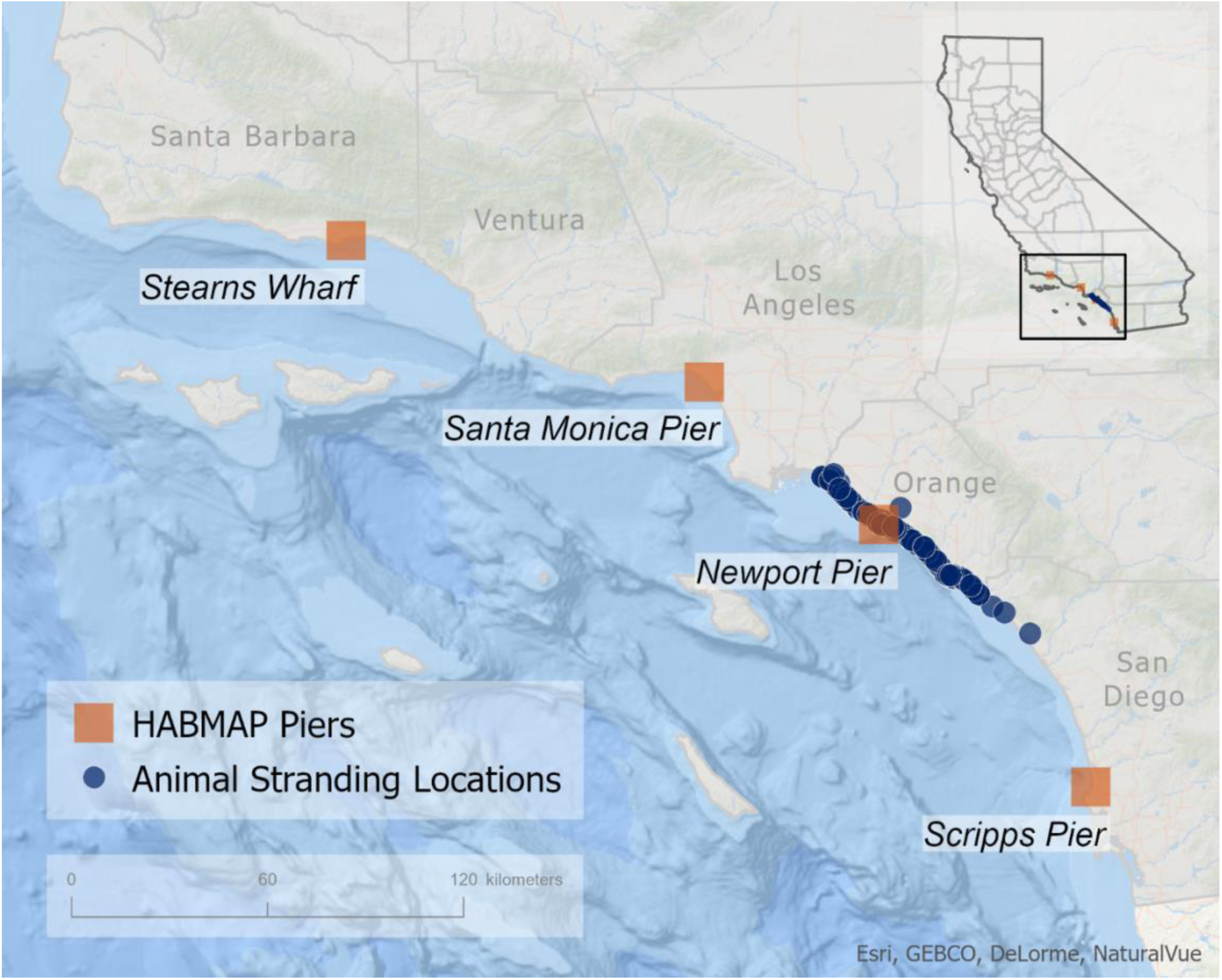
Map of regional domoic acid monitoring stations and California sea lion stranding locations. Location of adult and subadult California sea lion stranding rescued between 2015 and 2019 in relation to weekly pier monitoring stations. Stranding locations are shown in blue circles and the four weekly HABMAP pier monitoring stations are shown with orange squares.

In 2017, offshore water sampling was conducted in response to signs of a region-wide domoic acid producing bloom at the pier monitoring locations and reports of California sea lions stranding with domoic acid intoxication clinical signs. Surface water samples were collected opportunistically in Santa Monica Bay on April 13 and April 17, 2017 and in the Santa Barbara Channel on May 22, May 25 and June 6, 2017. Samples were also collected on the San Pedro Shelf region on April 18 and 19, and May 10 and 11, 2017 from both surface waters and the chlorophyll maximum located in the subsurface waters (depths ∼20m). A total of 73 discrete pDA samples were collected and analyzed as a part of these 2017 offshore sampling efforts following the same method applied to the pier monitoring samples.

### Marine Mammal Stranding Rates and Clinical Diagnoses

Pacific Marine Mammal Center (PMMC) is a member of the West Coast Marine Mammal Stranding Network, which was established under the United States Marine Mammal Protection Act by the National Oceanic and Atmospheric Administration (NOAA) to respond to stranded marine mammals. PMMC rescues, rehabilitates, and releases sick and injured marine mammals along the Orange County coastline, which is approximately 68 km of the Southern California coastline (**Figure 1**). All marine mammal rescue and rehabilitation activities are conducted by PMMC under a Stranding Agreement with National Marine Fisheries Service/NOAA.

Patient case records from 2015-2019 for California sea lions (hereafter, sea lions) were reviewed and collated for each animal’s case presentation including sex, age class, seizure activity, postictal signs, comatose, abortion, and whether the sea lion was diagnosed by the attending veterinarian with domoic acid intoxication following the behavioral diagnostic criteria outlined by Gulland et al., (2002). Multiple veterinarians made diagnoses during the time series based on these criteria. Age classes were defined following Greig et al. (2005) using dentition, straight length, pregnancy status, and sexual dimorphism to classify a sea lion’s age class. The following categories included in this study: juvenile/subadult male, subadult female, adult male, and adult female. Younger sea lions in the pup and yearling age classes were excluded as sea lions within these age classes are generally the most common patients at the center and often strand due to malnutrition (Bejarano et al., 2008a), and rarely present with acute or chronic domoic acid intoxication during blooms. Adult and subadult age classes are the most commonly affected by domoic acid intoxication (Bejarano et al., 2008a). Only live sea lions that were rescued on the Orange County coastline, brought to PMMC’s facility, and assessed by the attending veterinarian were included. Adult and subadult sea lions that died on the beach were excluded from the study cohort because an attending veterinarian could not assess clinical signs to diagnosis domoic acid intoxication prior to death.

### Statistical Analyses

Generalized linear autoregressive moving average (GLARMA) models were used to identify whether domoic acid concentration at each of four piers best predicted combined adult and subadult sea lion strandings along the Orange County coastline. The GLARMA model approach was selected for this analysis since it is well suited for use with observational, count based time series data in which counts are relatively small, have non-normal data distributions, and display serial dependence (Dunsmuir, 2015). The concentrations of domoic acid measured at each pier monitoring location along with past stranding events (0-6 weeks prior) were explored as potential explanatory variables for the presence/absence of a stranding event. Cross correlation functions were used to identify if domoic acid concentrations observed at pier locations led or lagged stranding events (**Supplemental Figure 1**). Based on autocorrelation patterns in the data (**Supplemental Figure 2**), the six and one week moving average values of stranding events preceding each observation were incorporated into the model, so that effects of domoic acid exposure could be explored after subtracting out the autocorrelated signal. Several stranding event scenarios were considered in our modeling efforts to determine if different demographics within the stranding cases yielded similar relationships to pier locations. These stranding event scenarios included total strandings, strandings where the sea lion was diagnosed as a suspected domoic acid case based on the behavioral criteria described above, strandings where the sea lion experienced a seizure, and strandings of female sea lions. The sample size was too small to model females that aborted fetuses while in treatment. We first queried a combined model that explored the effects of domoic acid concentration at all four piers. We then dropped each pier with lowest predictive power, in sequence, from our model, to determine whether a model with data from fewer piers was more informative. Akaike’s information criterion (AIC) was used to compare the performance of each model with lower AIC values indicating better fit models than higher AIC values in each modeled stranding scenario. The best fitting model for the total strandings prediction scenario was validated using a forward prediction approach. These analyses were conducted using the glarma package in R (Dunsmuir and Scott, 2015).

While the GLARMA model identifies statistical associations accounting for autocorrelation in stranding patterns, we also wanted to identify the general relationship between pDA concentration and strandings, ignoring autocorrelation patterns, in order to identify pDA thresholds that were associated with elevated stranding probabilities. Currently, there are no existing water thresholds with which to evaluate monitoring data for an increased risk of ecosystem impacts. Therefore, Poisson Regression modeling was used to explore association between domoic acid concentrations and adult sea lion stranding events ignoring temporal autocorrelation of time. This analysis was conducted using observations from the most strongly explanatory pier location, Stearns Wharf, which had been identified and previously found to be statistically significant in all of our modeling scenarios, using the GLARMA approach described above. Models were fit to identify the relationship between pDA concentration and the probability that at least one, or at least two sea lion stranding events occurred in a given week. This general linear model was conducted using the glm function, with Poisson family using the ‘stats’ package in R. Probabilities of one and two stranding events were generated by converting the lambda (λ) parameter generated by R’s prediction function into probabilities of more than one and more than two strandings using the ‘dpois’ function in the ‘stats’ package in R.

## Results

### Pier Based and Offshore Domoic Acid Dynamics

Detectable pDA concentrations were observed within the Southern California region each year between 2015 and 2019 (**Figure 2C-F**), with a significant domoic acid producing bloom of *Pseudo-nitzschia* was observed throughout the region in the spring of 2017. Detectable concentrations of domoic acid were observed at all pier locations routinely between March and early June, though concentrations varied substantially across piers during this event. Santa Monica Pier and Stearns Wharf had the highest observed domoic acid concentrations with multiple observations of pDA >1 μg/L, Newport Pier and Scripps Pier had lower concentrations of pDA that were generally <1 μg/L except for one observation in the first week of May at Scripps Pier of 2.07 μg/L. The highest concentration of domoic acid observed across pier stations peaked at 14.4 μg/L at Santa Monica Pier (**Figure 2D**). Offshore observations of pDA were more irregular than those at the piers, but these observations generally showed some marked spatial differences in pDA concentrations (**Figure 3)**. The highest offshore concentrations were detected offshore of Stearns Wharf in the Santa Barbara Channel (17 μg/L) and offshore of Newport Pier on the San Pedro Shelf (10 μg/L). Offshore and subsurface concentrations were often higher, sometimes by an order of magnitude or more, than those observed at Newport Pier (**Figure 3C**). Offshore observations in the Santa Barbara Channel and Santa Monica Bay generally showed less of a distinct difference from the pier observations (**Figure 3A-B**), but these observations were limited to surface grabs and were patchier than observations offshore of Newport Pier (**Figure 3C**).

**Figure 2.**
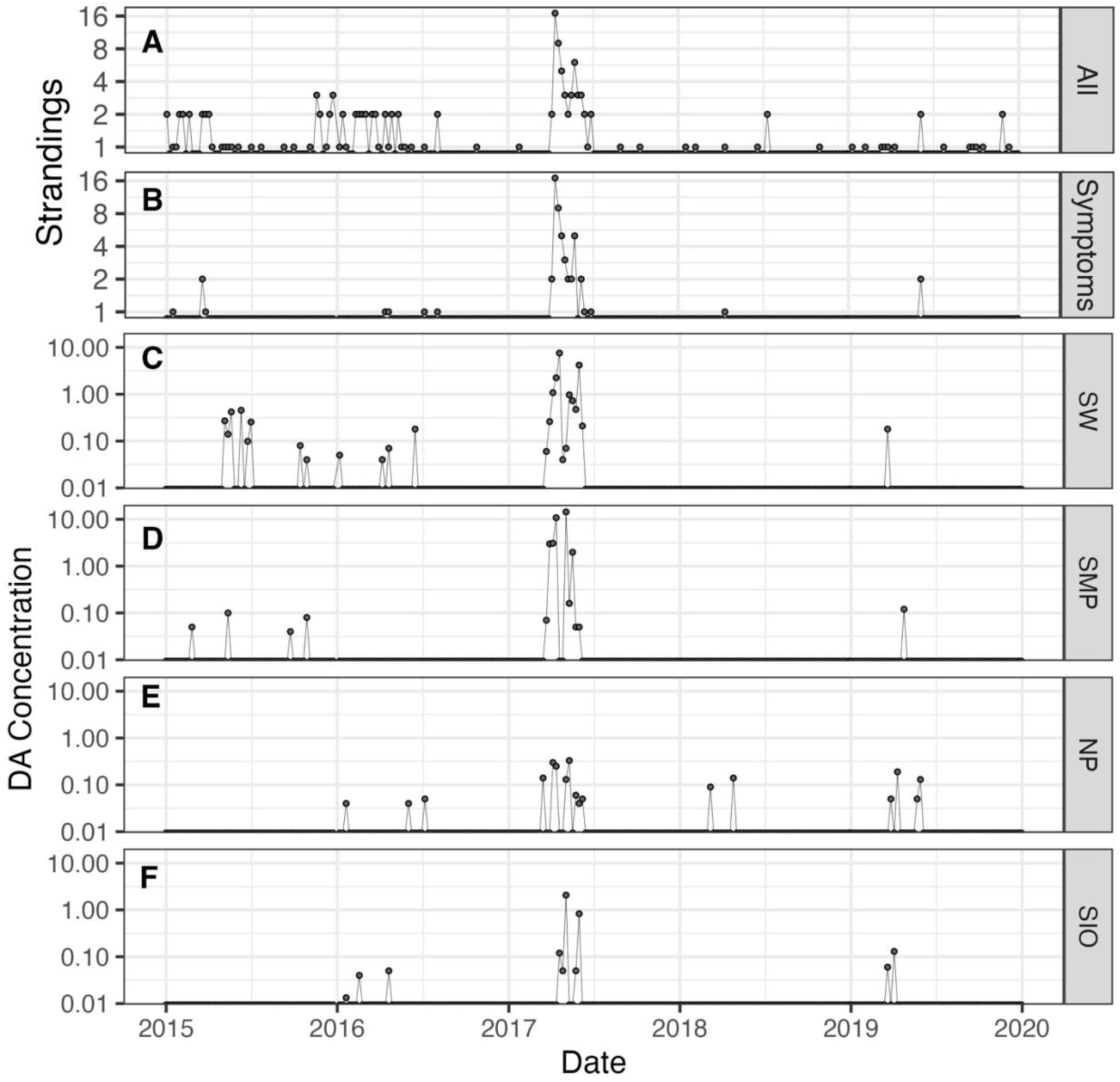
Time series of sea lion stranding counts and domoic acid concentrations from 2015 to 2019. Time series of total adult and subadult California sea lions (A), stranding patients that exhibited domoic acid symptoms while in treatment at PMMC (B) and particulate domoic acid concentrations (μg/L) at Stearns Wharf (C), Santa Monica Pier (D), Newport Pier (E) and Scripps Pier (F). Domoic acid concentrations are plotted on a semi-log scale.

**Figure 3.**
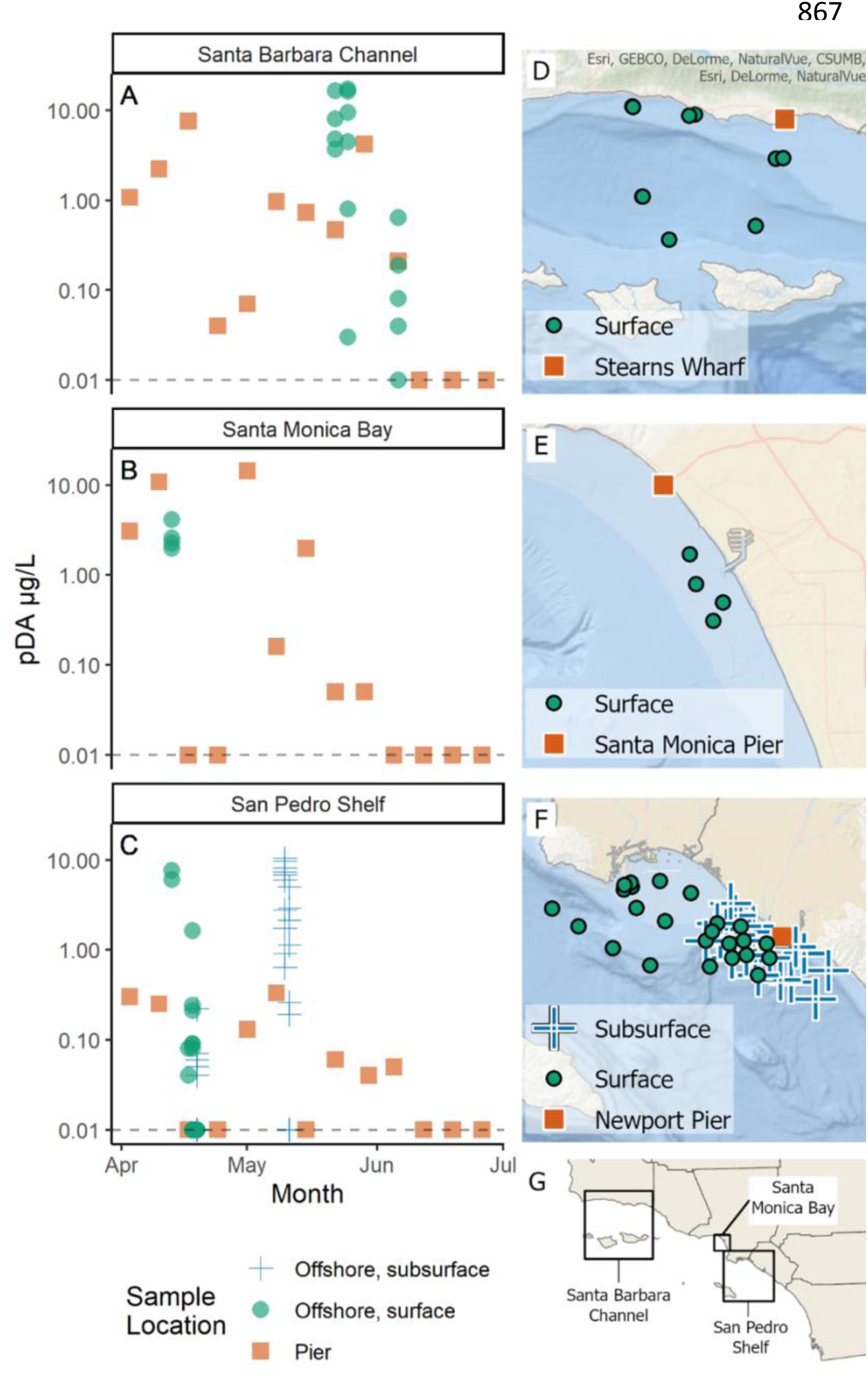
Pier based and offshore domoic acid concentrations observed during a large bloom event in 2017. Particulate domoic acid concentration (A-C) and location where samples were collected (D-G) during the 2017 opportunistic event response survey (green circles indicate surface samples and blue crosses indicate subsurface samples) in relation to routine pier monitoring (orange squares). Domoic acid concentrations are plotted on a semi-log scale with observations below the detection limit intersecting the dashed line at 0.02 μg/L. Geographic extent of three regions of interest is shown in G.

Observations of detectable domoic acid were more infrequent in the periods preceding and following the large bloom in 2017. In 2015, pDA was only detected at Stearns Wharf and Santa Monica Pier monitoring sites, with toxin detections ranging between 0.04 μg/L to 0.45 μg/L (**Figure 2C, D**). There was a period of fairly consistent detection of pDA between May and early July at Stearns Wharf, which was representative of a historically widespread bloom of *Pseudo-nitzschia,* which occurred across the U.S. West Coast (McCabe et al., 2016). However, this event was not observed at the other three piers to the south. In 2016, domoic acid was observed intermittently at Stearns Wharf, Newport Pier and Scripps Pier in the first half of the year at concentrations <0.50 μg/L. Following the 2017 bloom, domoic acid was least commonly observed at the pier locations in 2018, with only two instances of domoic acid detection at concentrations of <0.15 μg/L at Newport Pier. Domoic acid was detected at all four pier locations in the spring of 2019, although concentrations were all below 0.20 μg/L with the highest concentrations observed at Newport Pier of 0.19 μg/L (**Figure 2E**).

### Annual sea lion stranding rates and dispositions

The Pacific Marine Mammal Center rescued and administered veterinary care to a total of 153 adult and subadult sea lions between 2015 and 2019 (**Figure 2A**). During this period, adult and subadult strandings generally comprised a minor percentage of the overall sea lions rescued each year ranging from 7% (38/528) to 12% (16/138) of sea lion cases, apart from 2017 where 54% (61/114) of total strandings were in this age demographic. Of the subadult and adult sea lions stranded in 2017, 80% (49/61) were diagnosed as suspected domoic acid cases. Temporally, 95% (58/61) of the sea lion strandings during this year coincided with when elevated domoic acid concentrations were observed in the environment with 56% (34/61) of the sea lions strandings occurring in April, 23% (14/61) in May, and 16% (10/61) in June. Most of the adult and subadult sea lions that stranded during this period displayed various clinical signs consistent with domoic acid intoxication, ranging from postictal behavior (obtunded to comatose), seizure activity, and abortion. Additionally, 97% (56/58) of these stranded sea lions between April and June were females (**Supplemental Figure 3A**), 11 of which aborted a pup while in the rehabilitation hospital. All aborted pups were non-viable or stillborn, estimated to be 7 to 8 months in fetal gestation. Neurological or seizure activity was reported in 48% (28/58) of the stranded sea lions during the bloom, either on the beach or once back at PMMC.

In the years surrounding 2017 (2015, 2016, 2018, and 2019) PMMC responded to strandings of substantially fewer subadult and adult sea lions. Prior to the 2017 event, 38 subadult and adult sea lions were rescued in 2015, and 31 in in 2016. In the two years following the event, 7 subadult and adult sea lions were rescued in 2018, and 16 in 2019. Cases where sea lions were suspected domoic acid cases or observed to have seizures were also fewer in the years surrounding 2017. Of the total subadult and adult sea lion cases annually, 11% (4/38) were suspected domoic acid cases in 2015, 13% (4/31) in 2016, 14% (1/7) in 2018 and 13% (2/16) in 2019, compared to 80% of cases in 2017 (**Figure 2B**). Seizures followed a similar pattern in 2015 and 2016 and were observed in 13% (5/38) of cases in 2015 and 19% (6/31) of cases in 2016. Interestingly, a larger percentage of sea lions had reported neurological or seizure activity in 2018 and 2019, occurring in 43% (3/7) and 38% (6/16) of cases, respectively (**Supplemental Figure 3B**). These cases may represent chronic domoic acid patients, versus acute domoic acid intoxication associated with smaller, more ephemeral domoic acid events.

### Spatiotemporal relationships between domoic acid observations and sea lion strandings

Total adult and subadult sea lion stranding events typically occurred in temporal clusters with multiple strandings occurring over a period of weeks along the Orange County coastline, followed by periods without strandings of adult and subadult sea lions (**Figure 2A, B**). This dynamic was particularly apparent in 2017 during the domoic acid producing bloom of *Pseudo-nitzschia* that extended across most of the region. During this period, more than 50 sea lions stranded within a span of 11 weeks. Autocorrelation, but not partial autocorrelation, was observed in the overall stranding time series (**Supplemental Figure 2**), indicating that stranding in a given week is positively associated to previous stranding rates, with local autocorrelation maxima at one and six weeks.

Adult and subadult sea lion strandings on the Orange County coastline over the time series often preceded the observation of pDA at the piers. At all of the piers, except Newport Pier, the strongest observed cross correlations occurred with 1-3 week lead between pDA observations and total strandings, meaning that increased strandings were typically observed prior to an increase of pDA at the piers (**Supplemental Figure 1**). However, at all piers but Scripps Pier, the second most significant cross correlation observed was with no lag in pDA concentrations. Since our goal was to be able to estimate the risk of sea lion strandings with the pDA observations collected at the piers, and lagged values of pDA were not among the top cross correlations, we did not implement GLARMA models that considered previous values of pDA concentration at any pier. Instead, we used a GLARMA model that considered one-and six-week moving averages of stranding counts as inputs to account for these relationships. Log ratio and Wald tests confirmed that the model is better represented by processes that consider time lags in sea lion stranding counts than ones that did not (Wald = 26.01, p < 0.01).

The GLARMA model, fit to total strandings, was tested with pDA observations from all four piers within the region. Comparing AIC values of different models suggested that the total strandings scenario was most accurately predicted from models that include three piers (**Supplemental Table 1**). The best model fit included pDA data from Stearns Wharf, Santa Monica Pier and Newport Pier, and the statistical significance denoted by the p-values, which decreased in significance from north to south. Interestingly, only pDA concentrations at Stearns Wharf and Santa Monica Pier were statistically significant (p < 0.05, **Table 1**), while Newport Pier observations improved the overall model fit (as measured by AIC values, **Supplemental Table 1**) but its coefficient was not statistically significant (p = 0.088; **Table 1**).

**Table 1:**
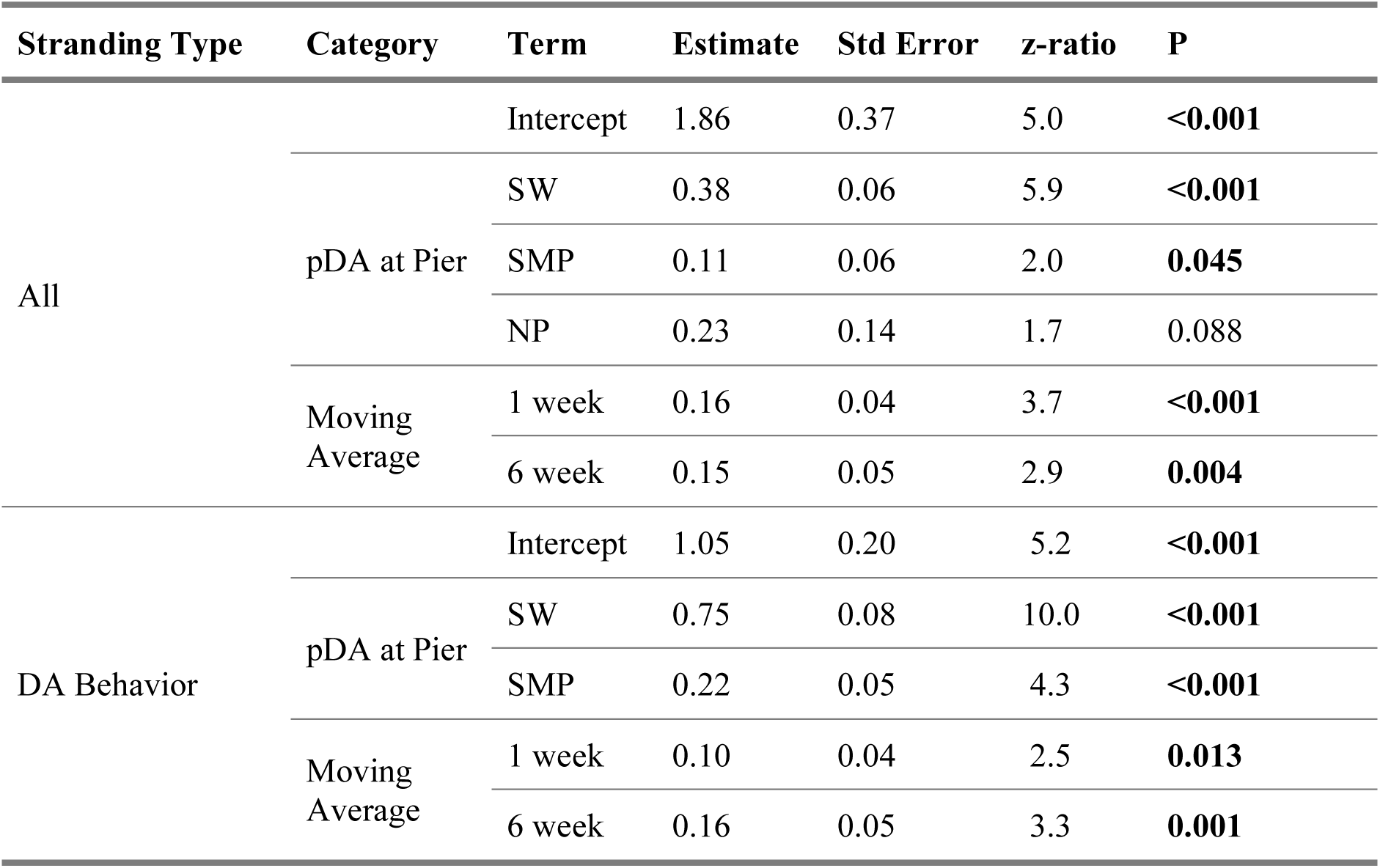
Model statistics for the total adult and subadult sea lions stranding scenario (Total) and the sea lions with domoic acid intoxication symptoms scenario. P-values < 0.05 are considered significant. pDA = particulate domoic acid.

Strandings in the other three scenarios considered also generally appeared in temporal clusters, and models were best fit using multiple piers (**Figure 2B, Supplemental Figure 2**). The GLARMA model for stranded sea lions that presented with domoic acid symptoms, incorporating one- and six-week stranding lag data, demonstrated the best fit when data from both Stearns Wharf and Santa Monica Pier were included as predictors (**Supplemental Table 1**). The inclusion of pDA data from the other two pier locations weakened model performance based on the AIC values (**Supplemental Table 1**). Sea lions exhibiting seizures and stranded females were modeled separately (**Supplemental Table 2**). All model scenarios were better fit by GLARMA processes than by equivalent general linear models (Wald Tests for each p<0.01).

The best fitting models for each sea lion stranding scenario were validated using a forward prediction approach, in which for each week models were re-fit on all data prior to that week and then strandings were fit using that week’s data and the one and six-week moving averages of strandings. Overall, the model for total stranded sea lions scenario performed well at predicting sea lion strandings from previous strandings and contemporaneous pDA concentrations from relevant pier monitoring locations (**Figure 4**). Predicted stranding probabilities mirrored observed strandings, as well as fitted stranding probabilities that used all of the data (including future data), for most months in the data set starting after the first year. Prior to 2016, there were not enough data to allow for model convergence. Furthermore, the model was “fooled” at the beginning of the domoic acid event in 2017 due to the negative lag term in the models built before 2017, resulting in substantially under-predicting the probability of additional strandings initially. However, from the second half of this event and forward, the model appeared to correctly predict the rest of the dataset. Forward predictions of the other three stranding scenarios (sea lions with symptoms of domoic acid exposure, subadult/adult female sea lions, and sea lions with seizures) showed less ability to predict stranding events (**Supplemental Figure 4**). The model that predicted stranded sea lions exhibiting seizures, however, only appeared to identify the peak in 2018. While it identified a non-zero baseline level of strandings, there did not appear to be enough cases of seizures outside of the 2017 stranding peak for the model to effectively identify elevated levels of seizure-strandings outside of this event (**Supplemental Figure 4**).

**Figure 4.**
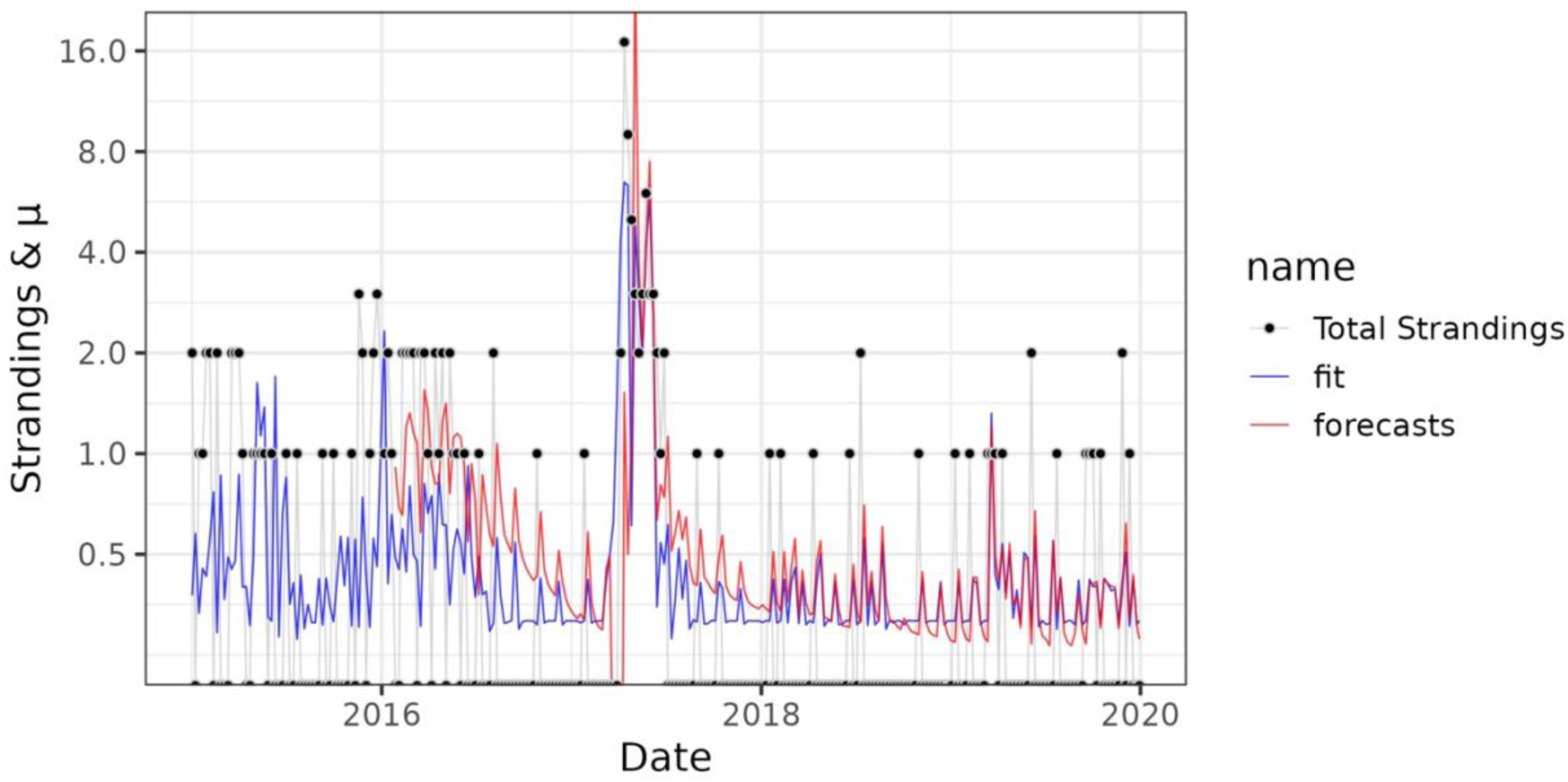
Modeled and actual observations of total sea lion strandings between 2015 and 2019. GLARMA forecast model of total strandings for study period. The y-axis corresponds to observed strandings and μ the predicted Poisson lambda value of the GLARMA model, corresponding roughly to the forecasted number and (if <1) probability of strandings. The x-axis corresponds to time. Black dots are observed stranding events. The blue line is the GLARMA model fit to the training data and forecasts are shown in red. The red line begins in 2016 because forecast models based only on data before 2016 fail to converge. The GLARMA forecast model is trained only with data preceding a given time point, and the domoic acid concentrations from that week. It then predicts the number of strandings for that week. The red line begins in 2016 because forecast models based only on data before 2016 fail to converge.

To better define the degree to which pDA observations are linked to negative ecosystem impacts, a simple model comparing pDA concentration at Stearns Wharf to strandings was also developed. This model, which ignored the temporal autocorrelation relationships described above, indicated that the probability of adult and subadult strandings events could be inferred from observed pDA concentrations (**Figure 5, Supplemental Figure 5**). Baseline stranding probabilities were never zero in these models, even when pDA was below the methodological detection limit. The baseline stranding probability for an individual adult/subadult sea lion in the total strandings scenario (when DA was set to the detection threshold of 0.02 μg/L was 44% per week (95% confidence intervals (CI): 38% – 49%) while the baseline probability of two or more sea lions stranding was 12% per week (CI: 8% – 15%; **Figure 5A**). Stranding probabilities increased with increasing concentrations of pDA in the environment. At a concentration of 0.05 *`μ`*g/L the probability of a single sea lion stranding was 55% (CI: 49% – 60%). At a concentration of 0.25 μg/L the probability of a single sea lion stranding was 81% (CI: 75% – 86%) (**Figure 5**). Multi-sea lion stranding probabilities were >50% per week at pDA concentrations of 0.25 μg/L and >80% per week at 0.75 μg/L.

**Figure 5.**
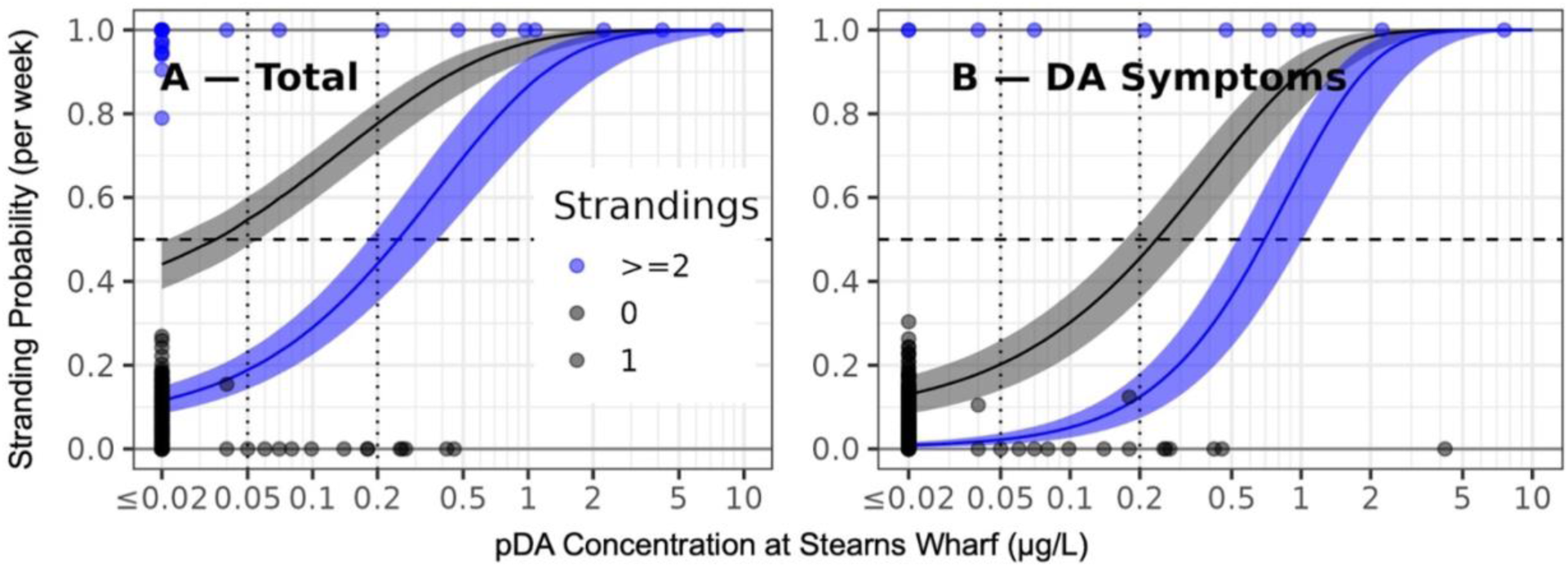
Probabilities of sea lion stranding events with increasing domoic acid concentrations. Observations and Poisson regression models of the relationship between domoic acid concentration and stranding scenarios, (A) total strandings and (B) sea lions exhibiting behavioral symptoms of domoic acid intoxication. Points indicate weekly observations. The x-axis corresponds to DA concentrations and the y-axis and point color reflect numbers of strandings. Points at or near the y-value of 0 correspond to weeks in which there were no strandings. Black points with y value at or near 1 correspond to weeks with one stranding. Blue points indicate weeks with two or more strandings. Black and blue bands indicate the probability, predicted by a Poisson regression model, of one or more (black) and to or more strandings (blue) at different domoic acid concentrations. Lines indicate the maximum likelihood probability and bands indicate two standard errors of that mean value.

## Discussion

All fitted models of adult and subadult sea lion strandings revealed a positive statistical link between strandings events and pDA concentrations at pier monitoring locations. Using the region-wide routine monitoring data, we were also able to identify a consistent correlation between strandings in Orange County and domoic acid at Stearns Wharf, nearly 200 km to the north of the stranding locations, across each of the stranding scenarios tested. Our best fitting models for each of the different stranding scenarios showed a pattern of decreasing significance from north to south across the region, underscoring the importance of considering the large geographic regions covered by sea lions when considering algal toxin exposure (Bernstein et al. 2021). Our results do not necessarily imply that domoic acid related marine mammal stranding events is solely driven by *Pseudo-nitzschia* blooms in the northernmost regions of Southern California, but in our time series, pDA observations at Stearns Wharf provide the best indication of when marine mammal strandings caused by domoic acid intoxication may occur. We believe this link could indicate that either pDA observations from Santa Barbara are potentially more representative than the other pier locations of the pDA concentrations in offshore and/or subsurface waters where sea lions forage, that sea lions do not strand near the areas where they became intoxicated, that sea lions throughout the region consume prey that originated near Santa Barbara, or a combination of factors.

### Spatial heterogeneity in Pseudo-nitzschia blooms complicates linkages between exposure and impacts

Our results suggest data from the nearshore pier locations can provide a reasonable approximation of when domoic acid related strandings are likely to occur, despite not fully capturing potential offshore and subsurface bloom dynamics. This finding is somewhat surprising in part because previous work has indicated that domoic acid observations collected at pier monitoring sites are often not fully representative of broader bloom spatial extent and magnitude within the region (Seegers et al., 2015; Smith et al., 2018). Estimates of remotely sensed chlorophyll-a have suggested a notable disconnect often exists between primary production patterns nearshore and those 2-4 km offshore in California (Frolov et al., 2013). Previous studies of blooms ecology within the Southern California region have shown that that there are sometimes significant differences between onshore *Pseudo-nitzschia* abundances and domoic acid levels compared to those both offshore and subsurface (Seegers et al., 2015; Smith et al., 2018). Notably, we found that subadult and adult sea lion strandings were typically observed prior to increased concentrations of pDA at most piers monitoring sites, limiting how much early warning these locations can provide for domoic acid related strandings.

Offshore observations collected during the large bloom event in 2017 highlight some notable spatial heterogeneity in domoic acid concentrations, particularly on the San Pedro Shelf region offshore of the PMMC stranding response zone and Newport Pier (**Figure 3**). While these observations do not have the same temporal resolution as the weekly observations collected at the piers, they highlight the potential for an order of magnitude difference in toxin concentrations between the nearshore and offshore regions, which may have significant implications for the predictive power of specific pier monitoring sites throughout the region. Interestingly, the difference between nearshore and offshore observations was not as pronounced in the Santa Monica Bay (**Figure 3B**) and Santa Barbara Channel (**Figure 3C**) regions, albeit observations were less extensive than in the San Pedro Shelf region and only represent a single major event (**Figure 3A**). Previous reports in the Santa Barbara Channel have demonstrated that there can be significant differences between concentrations of pDA at Stearns Wharf and those offshore in the Channel, with instances of offshore pDA concentrations exceeding those observed at pier stations by one or two orders of magnitude (Umhau et al., 2018). However, when considered over a five-year period between 2009 and 2013, there were not significant differences between concentrations of pDA routinely sampled offshore stations and those at Stearns Wharf (Umhau et al., 2018). This suggests that Stearns Wharf has good explanatory power because it is, on average, more representative of offshore domoic acid dynamics, which affect sea lions. Comparable pDA datasets to those presented in Umhau et al. (2018) do not currently exist offshore of Santa Monica Pier, Newport Pier, or Scripps Pier, therefore it is unclear how representative these locations are, on average, of more offshore conditions.

### Sea lion ecology results in variable domoic acid exposure

The life histories of sea lions contributes significantly to their domoic acid exposure and resulting stranding dynamics, particularly for acute exposure scenarios (Bargu et al., 2010). The main breeding colonies along the U.S. West Coast for sea lions are located on the Channel Islands, including San Miguel, San Nicolas, Santa Barbara and San Clemente Islands (Lowry and Forney, 2005). San Miguel, which is about 75 km offshore of Stearns Wharf and the most northern of the island chain, is the largest of the sea lion rookeries in the region (Le Boeuf and Bonnell, 1980; Lowry et al., 2021). The distribution of sea lions along the coast varies based on age and sex (Melin et al., 2000; Bargu et al., 2010). Our study focused on subadult and adult sea lions, which are either on the onset of sexual maturity or are sexually mature (Sinai et al., 2014). The breeding season occurs between May and August, during which time sexually mature males and females reside on the rookeries. Males rarely leave the islands to forage during the breeding season in order to maintain their territories (Bargu et al., 2010). Males spend the remainder of the year foraging along broad stretches of the U.S. West Coast as far north as Washington state (Weise et al., 2006). Female sea lions give birth between June and July, and typically remain with their offspring for the duration of their lactation period, which typically lasts for about one year (Gerber et al., 2010). During this period, these females split their time between nursing and foraging both offshore and on the continental shelf around the rookeries (Melin et al., 2000). Non-lactating females, however, disperse along the California coast following the breeding season (Melin et al., 2000). Domoic acid producing blooms exhibit a strong seasonality in Southern California, being most common in the spring and early summer (Smith et al., 2018), overlapping with the latter half of the maternal care period for lactating females and the beginning of the breeding season. Sea lions prey on several fish species known to accumulate domoic acid, including sardines and anchovies (Bernstein et al. 2021). Thus, the proximity of Stearns Wharf to the rookeries on the Channel Islands, and San Miguel in particular, may contribute to the consistent significance of the pDA observations at this location in our different stranding scenario models since a large proportion of the sea lion population reside and feed in that area when bloom events are common in in Southern California.

Most of the adult and subadult sea lions rescued by PMMC during the study period with symptoms of domoic acid intoxication were female, highlighting that bloom timing and sea lion ecology influences stranding demographics. This was particularly apparent in 2017 when a significant bloom event occurred. This event was first detected in April and ended in June, overlapping with the period when many of the region’s female sea lions would be foraging around the Channel Islands and surrounding regions. These observations are consistent with a stranding demographics study that indicated that adult females were between 47%-82% of the domoic acid cases reported by members of the California Marine Mammal Stranding Network between 1998-2006 (Bejarano et al., 2008a). Bargu et al., (2010) has previously highlighted how stranding demographics in Monterey Bay, CA may be related to temporal bloom dynamics. A bloom in the Monterey Bay region in March and April of 2007 was linked to the stranding of adult and subadult male sea lions, who were likely exposed to the bloom as they migrated though the area from the north to the breeding sites in the south (Bargu et al., 2010). The timing of blooms may therefore result in stronger impacts on specific demographics of the population, which has important population level effects in marine mammals that are currently understudied (Bejarano et al., 2008b).

### Study Caveats

The present study focused on the stranding patterns of adult and subadult sea lions, which likely contributed to the clearer relationship with domoic acid concentrations in the water than in previous studies. The causes of sea lion strandings are multiple, with some of the most commonly reported causes along the California coast being malnutrition, leptospirosis infections, and trauma (Greig et al., 2005). Yearlings are typically the most numerous age class encountered by stranding networks and they most often strand due to malnutrition (Greig et al., 2005). Furthermore, the domoic acid intoxication cases in this study were identified based on published behavioral diagnostic criteria (Gulland et al., 2002; Goldstein et al., 2008) and analytical confirmation of the toxin in patient fluids was not conducted. Although analytical confirmation of domoic acid in animal fluids or tissues is ideal for confirming a diagnosis, the behavioral diagnostic criteria, particularly for acute cases, are quite distinctive from many of the other leading causes of stranding in older sea lions. Thus, by excluding younger age class sea lions for our analyses, we were able to minimize the inclusion of sea lions that stranded for reasons other than domoic acid toxicosis and effectively model the relationship between domoic acid presence in the environment and related stranding events for multiple stranding scenarios.

The results of the present study are most applicable to linking environmental observations of domoic acid to stranding events related to acute intoxications. Acute intoxication events typically involve temporally clustered strandings, while sea lions presenting with signs of chronic domoic acid exposure often strand more sporadically and might be decoupled from significant bloom events (Goldstein et al., 2008). Our models all looked for temporal correlations between domoic acid in the environment and strandings and therefore, describe strandings related to acute, rather than chronic, exposures. When examining routine environmental observations of domoic acid and toxigenic *Pseudo-nitzschia* species from 2004-2007 in Monterey Bay, Bargu et al., (2012) reported a weak relationship between stranded sea lions of all age classes with signs of both acute and chronic domoic acid exposure. The relationship was improved, however, when only sea lions with acute symptomatology were included in the analysis (Bargu et al., 2012). Goldstein et al., (2008) reported a four-month time lag between sea lions that stranded with signs of acute domoic acid exposure and those that stranded with signs of chronic exposure along the central and northern coast of California. These observations all point to the decoupling between bloom dynamics and sea lions that strand with chronic or low level domoic acid exposure, making these types of strandings much harder to model and predict.

### Statistical considerations

Sea lion stranding data are challenging to analyze using many conventional time series approaches because they are autocorrelated count data. While ARIMA approaches (Shumway and Stoffer, 2010; Box et al., 2015), including general additive mixed models with autocorrelation functions (Cram et al., 2015) are conventional for time series data, these approaches expect the dependent variable to be normally distributed and so are not suitable for our count data, which are better modeled with a Poisson distribution. Similarly, while Poisson regression approaches can be used for count data (James et al., 2013), and while general additive models that assume Poisson or negative binomial distributed data are common for count data (Zuur et al., 2009), these approaches do not account for autocorrelated data. Hierarchical Bayesian approaches offer a possible solution to this problem (Tunaru 2002), though we note that the ability to work with autocorrelated Poisson data was removed from the commonly used BRMS package (Buerkner 2016), so implementation of such an approach likely requires additional validation. Therefore, we found the GLARMA approach was a good model for this data and anticipate that it will continue to be useful for future time series of sea lion strandings and similar phenomena.

One major goal of this study was to identify pDA concentrations of concern by identifying pDA levels that are associated with elevated stranding probabilities. A challenge of using GLARMA, and indeed any autocorrelation approach is that the modeled probability is dependent not only on observed domoic acid concentrations, but also on previous occurrences of strandings and so model outputs are all contingent on previous stranding levels and preventing determination of thresholds of concern. Therefore, to identify such a threshold, we used a simple Poisson regression model to determine the probabilities of one or two strandings, at different domoic acid concentrations, ignoring previous stranding levels. This simpler modeling approach was reasonable given that the GLARMA models, along with multiple mechanistic and field studies, have demonstrated associations between domoic acid and strandings that were not purely due to autocorrelation. Ultimately, this allowed for the identification of potential pDA levels of concern independently of summarizing recent stranding histories, which increases the utility of resulting numbers for risk assessment.

### Conclusions and Future Directions

The spatial and temporal heterogeneity in domoic acid producing blooms and sea lion habitat usage has previously made it difficult to link environmental observations to specific ecosystem impacts. Developing more quantitative linkages between environmental algal toxin concentrations and ecosystem effects is beneficial for both HAB ecologists and marine mammal stranding networks. The present study focused on a five-year period between 2015 and 2019, during which only one major domoic acid related stranding event occurred. Our analysis was also limited to stranding events occurring along the Orange County coastline. Future studies should explore these relationships across longer timescales to better understand the additional influences of low frequency changes in the environment, such as shifting climate modes, which might cause variations in both bloom development, marine mammal behavior and domoic acid intoxication patterns. Furthermore, relationships between environmental concentrations of domoic acid and strandings over broader geographic areas should also be examined, since there is potentially substantial variability in both bloom dynamics and localized marine mammal ecology over larger regions.

The relationship between offshore and subsurface blooms in the region and marine mammal stranding is not well studied due to the sparsity of routine offshore HAB observations. Domoic acid has been detected in several different marine mammal species stranded in Southern California that are known to forage in offshore pelagic and benthic habitats (Torres De La Riva et al., 2009; Fire et al., 2010). In some cases, marine mammals, including sea lions, strand and display signs of acute domoic acid exposure in advance of observations of toxigenic *Pseudo-nitzschia* cells in the nearshore environment (Torres De La Riva et al., 2009; Bargu et al., 2012; Broadwater et al., 2018), pointing to the occurrence of offshore events that precede nearshore bloom observations. Using cross correlation functions, we also observed evidence of sea lion strandings slightly leading observations of increased pDA at most pier locations in our time series (**Supplemental Figure 1**), which could be attributed to sea lions being exposed to domoic acid related to offshore and/or subsurface blooms that were not yet detected at the pier locations. More routine observations of offshore and subsurface bloom dynamics regionally could help better resolve their connectivity (or lack thereof) to nearshore bloom dynamics and would likely improve the relationship and potential predictive power of future forecasting efforts. Until these dynamics are better resolved, pairing the routine pier monitoring observations with monitoring of sea lions strandings in Southern California is useful in directing when it is necessary to perform offshore and subsurface monitoring for domoic acid to facilitate more rapid identification of these harder to detect bloom events.

We identified pDA concentrations of potential concern based on the probability that specific concentrations could result in ecosystem impacts, with 0.05 μg/L pDA and 0.25 μg/L pDA representing two potential levels of concern based on increasing risk of sea lion stranding events. Ultimately, developing these types of relationships can help to move towards the ability to forecast marine mammal stranding events related to HABs. The capacity to model and forecast domoic acid producing blooms is increasing; it would be beneficial to link these improved forecasts to ecosystem impacts, such as marine mammal strandings. Currently, the California coast has an operational forecasting model, California Harmful Algal Risk Mapping (C-HARM; Anderson et al., 2016). With additional research, C-HARM or similar model products could be expanded to also consider marine mammal stranding risks based on these pDA levels of concern, thereby linking the model outputs to ecosystem effects.

## Author Contributions

JS, JAC, VH, and ACD wrote the manuscript, MB, and DS contributed significantly to initial design and revisions.

JS, MB, VH, DS, and ACD conducted the data acquisition and dataset assembly, JS, JAC, and ACD analyzed and synthesized the data with significant contributions from DS.

All authors contributed significantly to the interpretation of the data.

All authors approved the submitted version for publication.

## Acknowledgements

The authors wish to thank the principal investigators, field staff and students associated with Cal-HABMAP pier monitoring efforts funded by the Southern California Coastal Ocean Observing System for collecting sustained observations of domoic acid since 2008. We also thank the animal care staff, volunteers, and donors at the Pacific Marine Mammal Center who assisted and supported rescue and rehabilitation efforts of the stranded marine mammals for the duration of this study. We wish to acknowledge the National Centers for Coastal Ocean Science (NCCOS) HAB Event Response Program for supporting offshore sampling efforts in 2017. We thank Meredith D. A. Howard, David A. Caron, Raphael M. Kudela, and Clarissa Anderson for coordinating event response sampling and analysis efforts in 2017. We also thank the Orange County Sanitation District, City of Los Angeles Environmental Monitoring Division, and the Santa Barbara Channel Keepers for offshore field sampling assistance. This document is NCCOSS HAB Event Response Publication #XX.

## Data accessibility statement

Harmful algal bloom observations made by Cal-HABMAP are publicly available on the program website: https://calhabmap.org/. California sea lion stranding records can be found at XXXX.

