## Supplemental Material for "Quantifying the linkages between California sea lion (*Zalophus californianus*) strandings and particulate domoic acid concentrations at piers across Southern California"

**Supplemental Table 1**. AIC values of models with different numbers of piers included. Lower AIC values indicate a better model fit within each stranding scenario tested. Piers are retained from north to south, so a model with only two piers for instance includes the Stearns Wharf and the Santa Monica Pier, but not the Newport or Scripps Piers. Similarly, the model with only one pier includes only the Stearns Wharf. The seizure model with four piers did not converge (DNC).

| **Stranding Type** | **Number of Piers** | **AIC** |
| --- | --- | --- |
| Total | 3 | 493.36 |
|  | 2 | 494.04 |
|  | 4 | 494.75 |
|  | 1 | 505.09 |
| DA Behavior | 2 | 206.14 |
|  | 4 | 206.36 |
|  | 3 | 206.37 |
|  | 1 | 223.76 |
| Female | 2 | 403.21 |
|  | 3 | 403.97 |
|  | 4 | 404.54 |
|  | 1 | 419.74 |
| Seizure | 3 | 229.10 |
|  | 2 | 229.33 |
|  | 1 | 235.53 |
|  | 4 | DNC |

**Supplemental Table 2:** Model statistics for the female sea lions stranding scenario and the sea lions with seizures scenario. P-values < 0.05 are considered significant. pDA = particulate domoic acid.

| **Stranding Type** | **Category** | **Term** | **Estimate** | **Std Error** | **z-ratio** | **P** |
| --- | --- | --- | --- | --- | --- | --- |
| Female |  | Intercept | 1.30 | 0.20 | 6.6 | **<0.001** |
|  | pDA at Pier | SW | 0.49 | 0.06 | 7.6 | **<0.001** |
|  |  | SMP | 0.19 | 0.04 | 4.4 | **<0.001** |
|  | Moving Average | 1 week | 0.15 | 0.04 | 3.7 | **<0.001** |
|  |  | 6 week | 0.18 | 0.06 | 3.0 | **0.002** |
| Seizure |  | Intercept | 1.608 | 0.5 | 3.10 | **0.002** |
|  | pDA at Pier | SW | 0.550 | 0.1 | 5.77 | **<0.001** |
|  |  | SMP | 0.119 | 0.1 | 0.91 | 0.364 |
|  |  | NP | 0.350 | 0.2 | 1.46 | 0.144 |
|  | Moving Average | 1 week | -0.202 | 0.1 | -1.95 | 0.051 |
|  |  | 6 week | 0.031 | 0.1 | 0.23 | 0.819 |

**Supplemental Figure 1:** Cross correlation function plots of total sea lion strandings (as the y-variable) along the Orange County Coastline versus particulate domoic acid observations (as the x-variable) at A) Stearns Wharf, B) Santa Monica Pier, C) Newport Pier and D) Scripps Pier. Relationships were observed on a weekly basis with each peak representing a forward or backward lag in weeks from zero. Blue dashed lines are 95% confidence intervals of an uncorrelated series and so lags that exceed these confidence intervals are statistically significant.

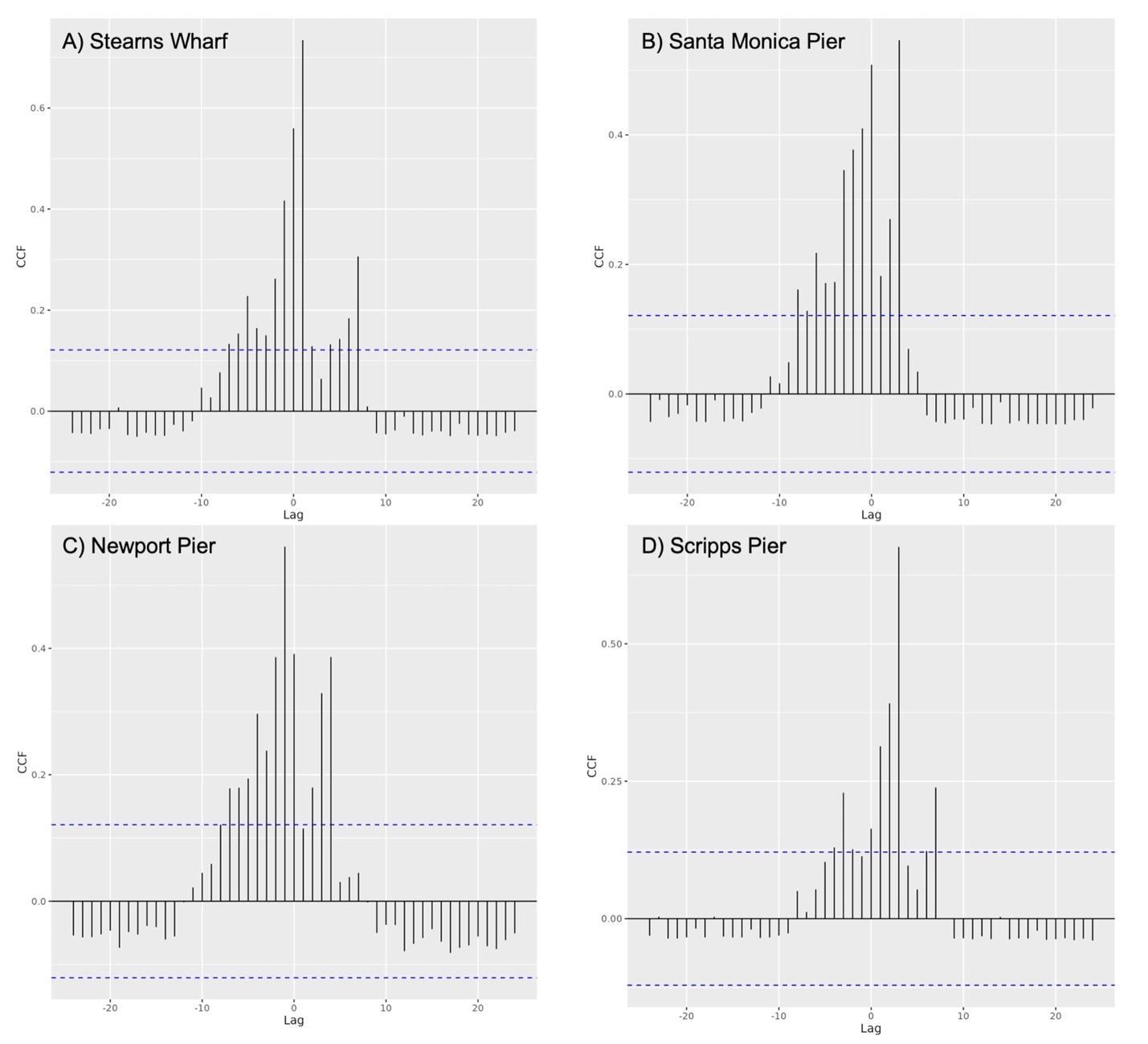

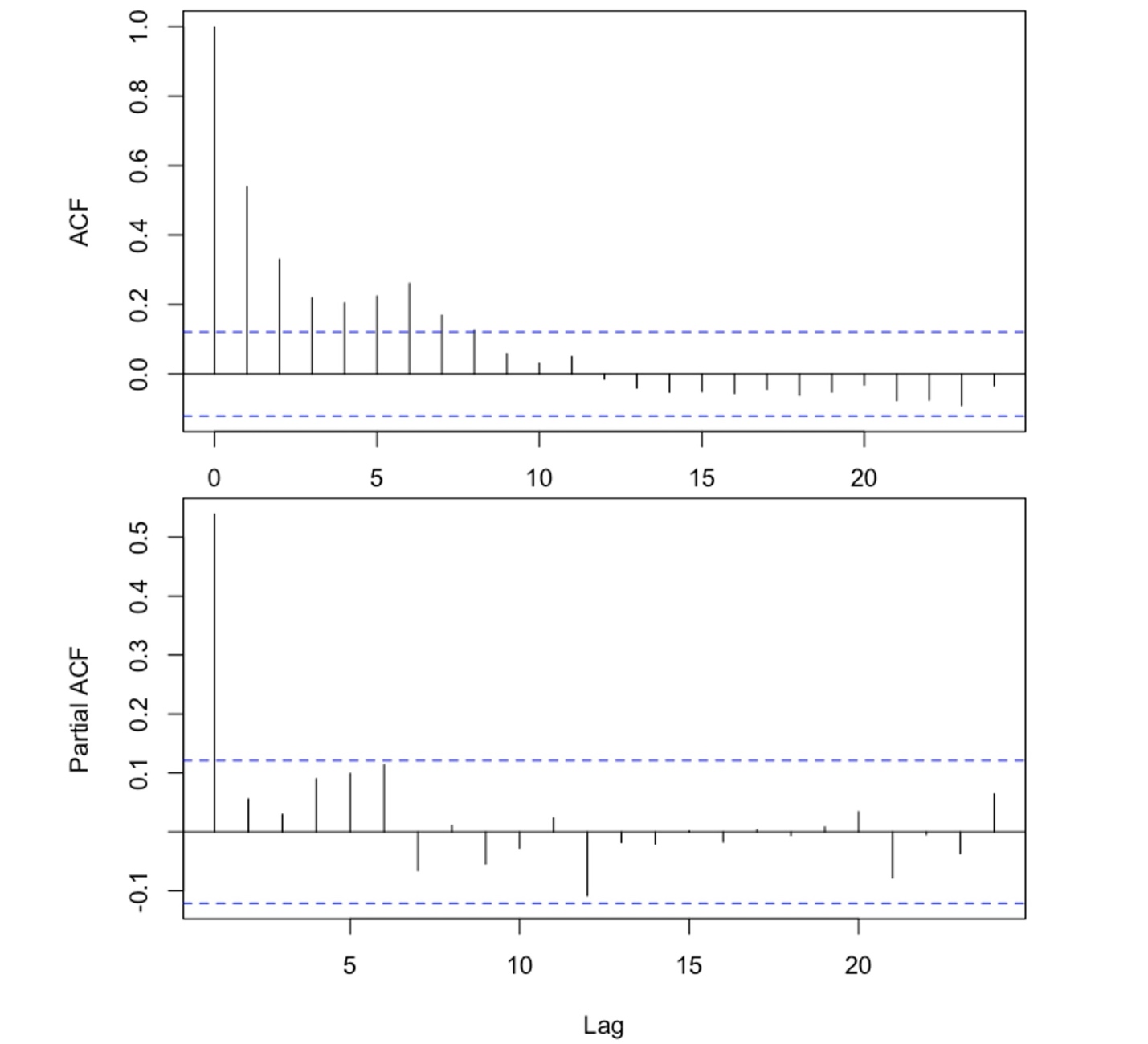
**Supplemental Figure 2.** Autocorrelation (AFC; A) and partial autocorrelation (PAFC; B) of the stranding data. Vertical bars indicate ACF and PACF scores (y-axis) corresponding to time lags of a given number of weeks (x-axis). Blue dashed lines are 95% confidence intervals of an uncorrelated series and so lags that exceed these confidence intervals are statistically significant.

**Supplemental Figure 3:** Time series of total adult and subadult female California sea lions (top), stranding patients that exhibited seizures while in treatment at PMMC (bottom).

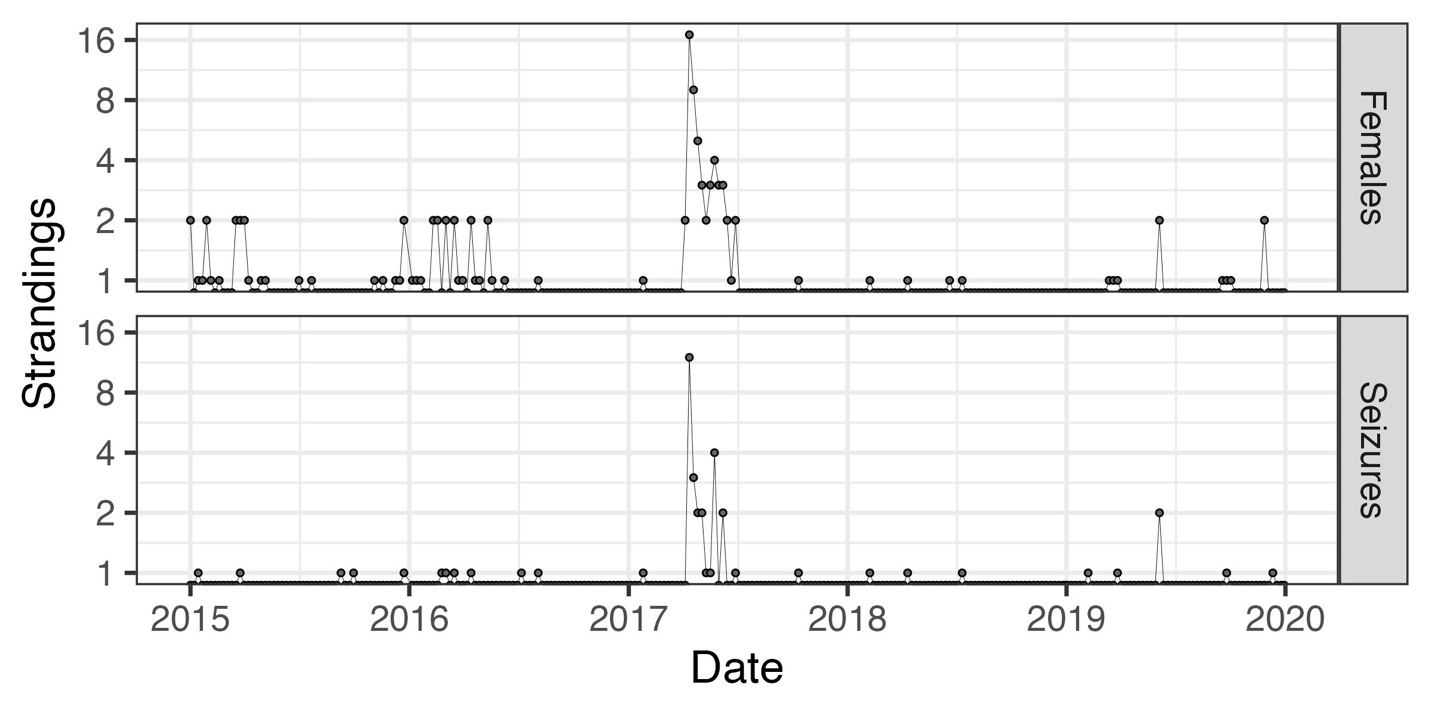

**Supplemental Figure 4:** GLARMA forecast model of total strandings compared to the other three stranding scenarios. The x-axis is the date. The y-axis corresponds to observed strandings and μ the predicted poisson lambda value of the GLARMA model, which corresponds roughly to the forecasted number and (if less than one) probability of strandings. Black dots are observed stranding events. The blue line is the GLARMA model fit to all of the training data. In the forecasts from the red line, the GLARMA forecast model is trained only with data preceding a given time point, and the DA concentrations from that week. It then predicts the number of strandings for that week. The red line begins in 2016 because forecast models based only on data before 2016 fail to converge.

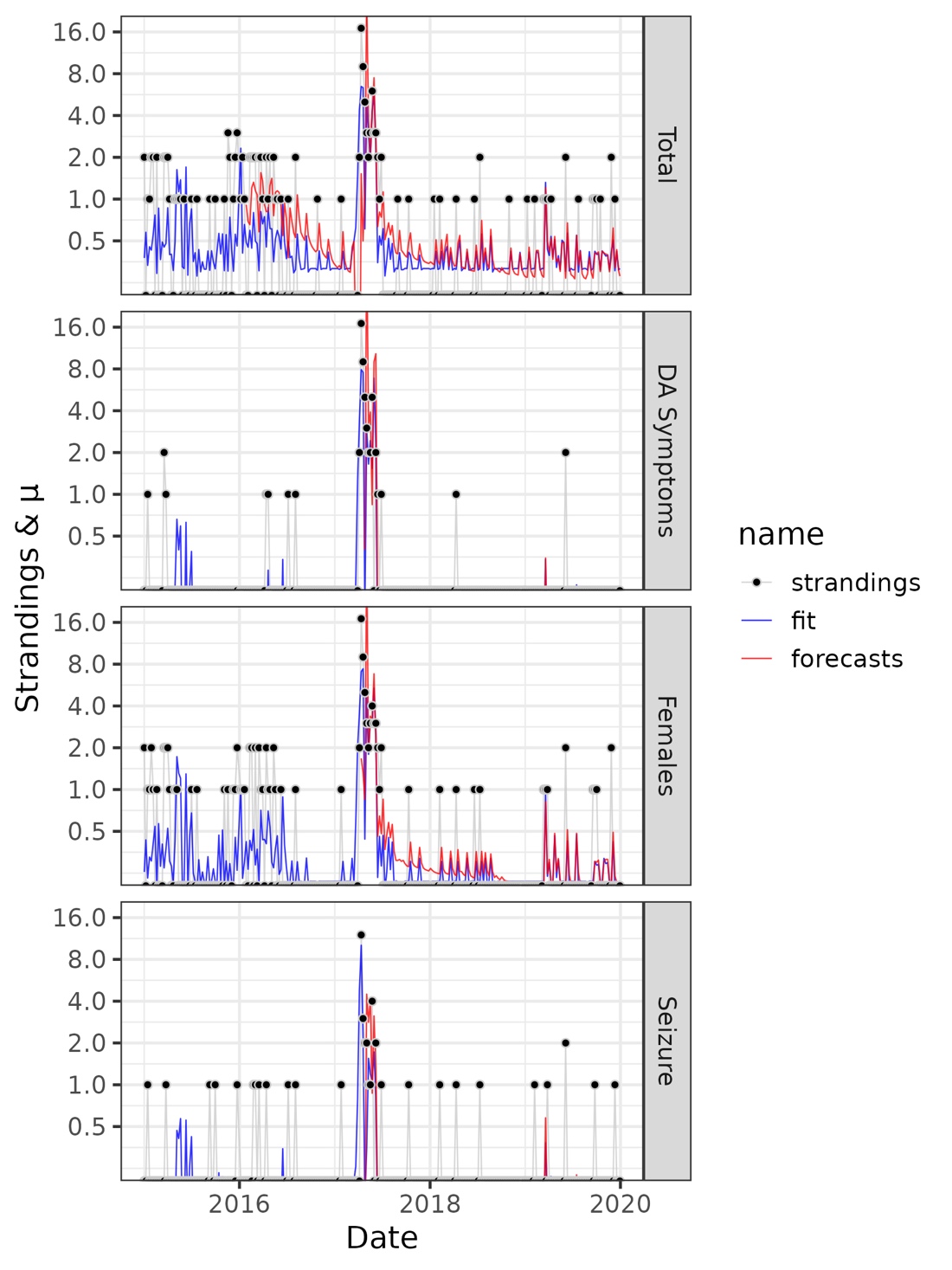

**Supplemental Figure 5:** Observations and Poisson regression models of the relationship between domoic acid concentration and stranding scenarios, (A) females and (B) sea lions exhibiting seizures while in treatement. Points indicate weekly observations. The x-axis corresponds to DA concentrations and the Y axis and point color reflect numbers of strandings. Points at or near the y-value of 0 correspond to weeks in which there were no strandings. Black points with y value at or near 1 correspond to weeks with one stranding. Blue points indicate weeks with two or more strandings. Black and blue bands indicate the probability, predicted by a Poisson regression model, of one or more (black) and to or more strandings (blue) at different domoic acid concentrations. Lines indicate the maximum likelihood probability and bands indicate two standard errors of that mean value.

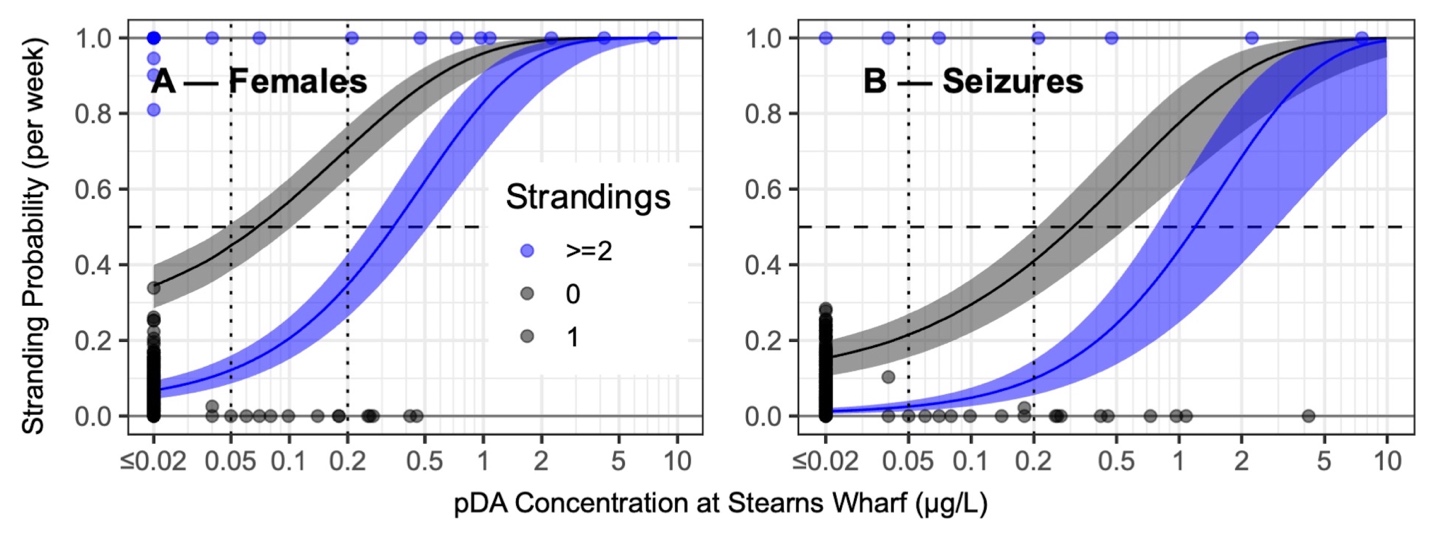
